# The simulation of functional heterogeneity in procedurally generated fibrotic atrial tissue

**DOI:** 10.1101/2022.11.28.518223

**Authors:** A.I. Kalinin, S.G. Kovalenko, A.K. Berezhnoy, M.M. Slotvitsky, S.A. Shcherbina, V.A. Syrovnev, V.A. Tsvelaya

## Abstract

The occurrence of atrial fibrillation (AF), one of the most common and socially significant arrhythmias, is associated with the presence of fibrosis sites. Fibrosis is the presence of non-conductive fibroblast cells, separating cardiomyocytes and introducing heterogeneity into the conducting atrial tissue. Thus fibrosis may be a substrate of spiral-wave reentry, provoking the occurrence of AF and is often associated with its persistent form. In this work, we propose for the first time a fundamentally new approach to modeling the fibrotic heart that takes into account the cellular structure of the tissue: a realistic texture of atrial tissue remodeled by fibroblasts is generated by the Potts model, and the local membrane potential of individual cells is calculated by the Courtemanche model. We have shown the occurrence of conductive pathways in such a system with a low proportion of fibroblasts (up to 10%) and revealed the connection of the form of the action potential (AP) of cells with their location in the tissue and the direction of the propagating wave front. The combination of these effects creates dynamic heterogeneity of the conducting tissue and affects the migration and pinning of spiral waves, which makes the model a potential tool for prognostic modeling of AP and search for ablation targets. The computer prediction of ablation targets (reentry nodes) will help to increase the efficiency of treatment of patients with persistent form of AF.

## 1. Introduction

Cardiovascular disease (CVD) is the leading cause of global mortality among working-age adults worldwide, with the proportion of CVD deaths increasing with increasing life expectancy. One of the most common cardiovascular diseases is atrial fibrillation (AF), which occurs in more than 1% of the total world population. Often a patient is first diagnosed with AF after suffering a stroke, which occurs in 25% of cases of AF. Other reason of social importance of AF - possibility of its transition to ventricular fibrillation, representing fatal danger [1]. Despite of high incidence of atrial fibrillation (AF) in the population, the question on its pathogenesis remains debatable, that is reflected in relative effectiveness of treatment of this disease from 50 to 80% in the one-year period [1–4]. Isolation of a pulmonary vein by means of catheter ablation can effectively treat some forms of AF, but recurrence rate remains unacceptably high (40-60 %) in patients with persistent AF (PsAF). A possible explanation of relative inefficiency of ablation is the fact that PsAF is often connected with atrial fibrosis, representing a substrate for which an arrhythmogenic predisposition zone extends beyond the ablative area. One of the leading hypotheses about the occurrence and maintenance of AF is the notion that a microreentry type excitation occurs in the pulmonary veins or in the atrial wall that has undergone partial fibrous remodeling. In different sources, this mechanism is also found under the name of rotor or driver activity [2,5–8].

In studies on cell cultures and computer models the conditions of microreentry emergence have been studied. It is believed that the main condition for the emergence and maintenance of reentry activity is the presence of non-conductive areas in the tissue - up to 70% fibrous tissue [9,10]. But in practice, the amount of fibrous tissue for the onset of AF is much less. It is known that cardiac tissue is significantly anisotropic, i.e. the velocity along the fibers in the heart can exceed the velocity across the fibers by more than two times in ventricles and ten times in atria [11]. The vast majority of modern mathematical models of cardiac tissue are based on the concept of cell syncytium, i.e. they proceed from the idea that internal volumes of cardiac cells are connected by special pores (connexons) [12]. However, this notion faces a number of contradictions between theory and experiment. It has been shown that on the subcellular scale, excitation propagation occurs in leaps and bounds - rapidly across the cell membrane and slowing down as it passes from cell to cell [12]. In addition, the duration of primary depolarization differs depending on the direction of wave propagation [13]. It is also known that channel blocking leads to disruption of anisotropy. Such effects can lead to the disturbance of symmetry of wave conduction in the heart and, consequently, to the occurrence of reentry [14,15]. Thus, the data on the conditions of emergence and stability of microreentry in human tissues are limited and are not reproduced fully on existing computer models.

One way to study fibrosis and anisotropy is through simulation modeling. Conducted in computer models for specific patients, such simulations can reconstruct atrial conduction based on late scans of the patient’s organ by magnetic resonance imaging (LGE-MRI) with gadolinium contrast, thus reconstructing a 3D atrial geometric model. Recently, such models have begun to appear to provide information on preservation and elimination of PsAF for patients with atrial fibrosis. For example, computer model OPTIMA [16, 17] creates an individual plan of ablation in the format compatible with clinical electro-anatomical navigation system (CARTO) [18]. But modern models have significant disadvantages. One of them is that they do not take into account cellular interactions, defining fibrosis as simply an area with medium reduced intercellular diffusion [9]. This analogy cannot give correct results because it does not take into account the discrete distribution of conductive and non-conductive cell types at the sites of fibrosis. That is, it cannot calculate microreentries on the fibrous substrate and determine whether they are the cause of recurrent AF [19] (in such a model, it is the border of the fibrous region that is arrhythmogenic). Therefore, these models cannot be used to predict the long-term effect of ablation and the probability of recurrence for each case of ablation. The present work supports the hypothesis of the fibrous substrate as the main arrhythmogenic zone, which determines the need to define new techniques for precise identification of ablation targets in the fibrous substrate in order to reduce the recurrence rate of AF.

The presented study shows that atrial fibrosis is a key link in arrhythmogenesis observed in patients with AF. The study of the morphological features of fibrosis led to the creation of a computer model of atrial fibrosis based on the identified relationships between fibrosis and the occurrence of arrhythmias, which predicts the conditions of AF. The model itself was first presented for atrial fibrosis in this work. It demonstrated the mechanism of microreentry formation and their steady state only by accounting for cell morphology and intercellular contacts. Subsequently, such model can lead to a prognostic model of patient’s atria to support decision making on treatment of AF, its early diagnosis and selection of a protocol of surgical ablation.

## 2. Methods and materials

### 2.1. Fibrous tissue generation, diffusion coefficient arrangement

The Potts model, whose Hamiltonian is formulated as follows, was used to generate the samples (formula 1):

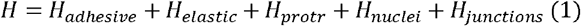

The formulation of the last term and the values of the constants are taken from [9], the other formulations are from [20]. Using Python code, the fixed coordinates were converted into diffusion D-values linking neighboring pixels. The sample required 50’000 MCS to generate, and the cell boundaries did not change significantly as the number of steps increased. The input data for the Potts model in this work are: the initial coordinates of fibroblasts (FBs) and cardiomyocites (CMs) that mixed randomly, the directions of parallel fibers specifying the tissue anisotropy.

### 2.2. Electrophysiological model

To calculate the membrane potential of atrial cardiomyocytes, we used the Courtemanche model, which is reduced to formulae (formula 2) and (formula 3):

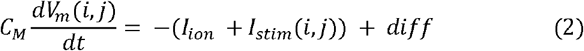

The formulations of the individual ionic currents and the values of the constants were taken from [21]. The constants from [22] were used to simulate the action potential (AP) of the tissue subjected to AF.

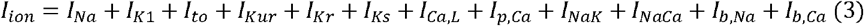

Formulas 2 and 3 are the same throughout for all pixels (i,j). Thus, the electrophysiology (conductivity of specific ion channels and their temporal characteristics) of each sample membrane element is the same.

### 2.3. Conductivity Simulation, Stimulation Protocols and Difference Diagram

The conduction simulation was implemented in Myokit (http://myokit.org/), the Courtemanche model formulation were imported from CellML Model Repository (https://models.cellml.org/cellml). Myokit was augmented with code (Python 3) that translates the image generated by the Potts model into the set of diffusion coefficients (D) used in the difference scheme and included in the last term of formula (2). This term is formed by the difference scheme, a detailed description of which is given in [23]. The time step value used was 1 ms, and the space step value was 2.5 μm.

For single stimulation, we applied a step pulse with an amplitude of 100 pA, localized in space (the area where I_stim_ is not equal to 0 is shown as a white rectangle in the figures). With the s1s2 protocol, the first stimulation is performed at the upper edge of the sample. The trigger for the second stimulation (square stimulation in the upper left quarter of the sample) is the return of V_m_ in the middle of the sample to a value of −70 mV.

### 2.4 Excitation propagation analysis (action potential morphology, activation maps)

The activation maps were constructed in Wolfram Mathematica, and the S-value was calculated using ImageJ to characterize the AP shape in each pixel:

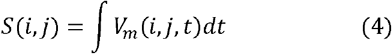

Diagramming and extraction of the local action potential form from the simulation was also performed using the ImageJ program.

## 3. Results

### 3.1. Formation of functional heterogeneity in fibrous tissue

To investigate the conduction properties of the sample, we generated a section of tissue with 10% fibrosis, and parallel directed fibers. The diffusion coefficients (D) in the difference scheme (see section 2.3.) were chosen so that the pulse propagation within one cell was as fast as possible, while the excitation transmission to the neighboring cell caused a delay. The model provides two mechanisms of impulse transmission: through gap contacts, the location of which is given by Potts’ model (formula 1) and due to the ephatic connection of cells (table 1, column 3). The second mechanism, which has not been previously implemented in the Potts model [9], may be an alternative to gap contacts [26] and has been shown experimentally [24] to play an important role in the formation of anisotropy [25] and the stability of conduction with a small number of gap contacts [26]. The conductivity of fibroblasts is equated to zero, which together with the difference scheme (section 2.3) describes the Neumann boundary condition (no-gap boundary).

**Table 1.** Boundary conditions in the Potts model for maintaining ephatic cell bonding.

| Membrane conductivity | Gap Junctions (GJ) conductivity | Alternative conductivity | Non-conductive areas |
| --- | --- | --- | --- |
| 551 $\mu\text{S}$ | 221 $\mu\text{S}$ | 10 $\mu\text{S}$ | 0 $\mu\text{S}$ |

Based on the conductance ratio and homogeneity of Formulas 2 and 3, we can assume that the shape of the action potential will be unchanged within each individual cell and may differ between neighboring cells (because cell size and total capacitance of the membrane bounding it vary, Figure 1). Figure 2 shows the passage of an excitation wave through a sample with 10% fibroblasts. The S-value is used to characterize the action potential shape (see Section 2.4.), Figure 2 demonstrates significant heterogeneity of the action potential within the sample, and the activation map shows the curvature of the leading front of the wave as it propagates through the sample. The front curvature cannot be explained by the boundary conditions alone (Table 1).

**Figure 1.**
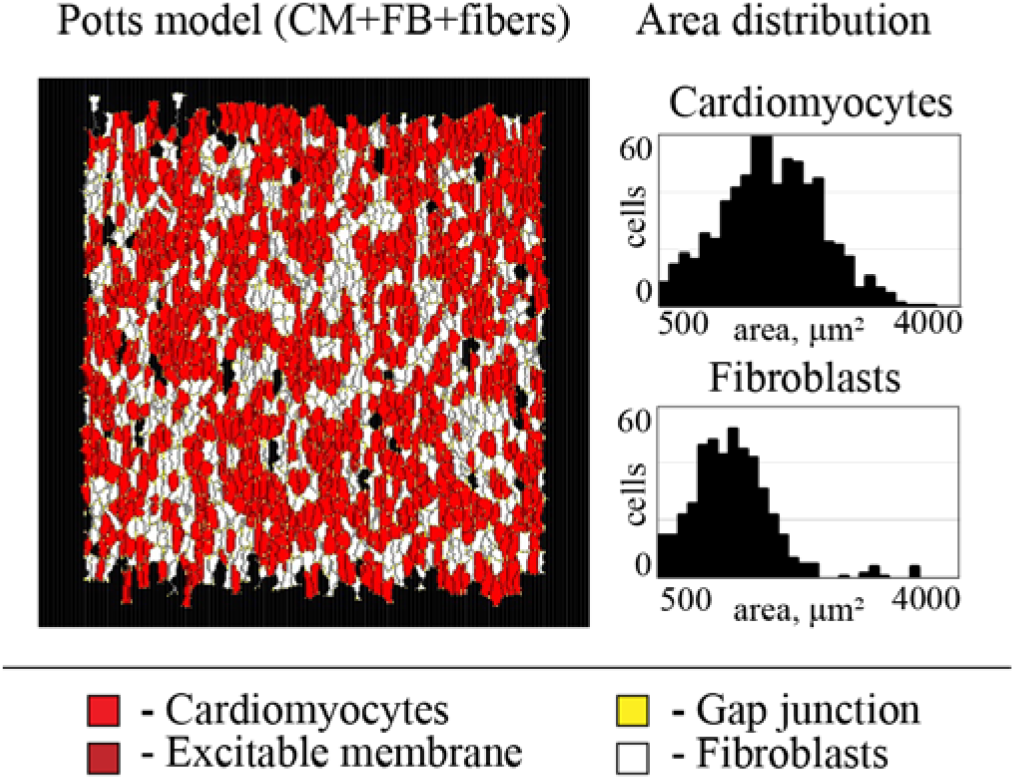
A sample of anisotropic atrial tissue with 50% fibroblasts, cardiomyocytes are shown in red, contacting membranes of cardiomyocytes in burgundy, and gap junctions in yellow. Fibroblasts (non-conductive cells) are shown in white. The diagrams on the right show the distribution of cells composing the specimen by occupied area.

**Figure 2.**
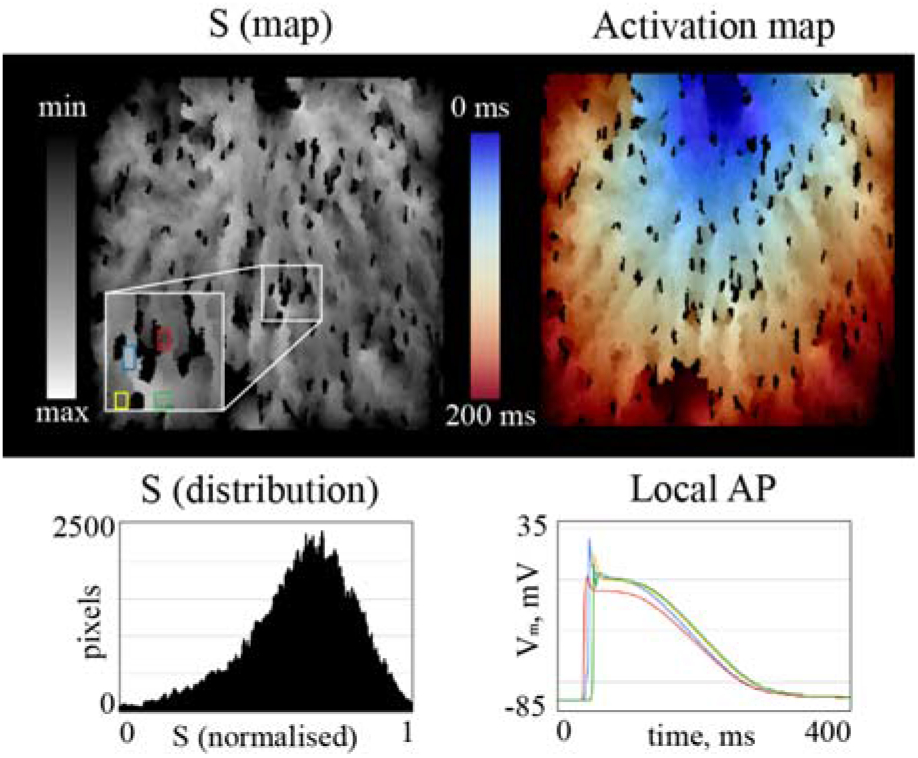
The map of the AP forms (left) and activation map (right) for the excitation wave triggered by the pulse in the center of the upper edge of the sample. The diagram on the bottom left shows the scatter of S-values in the sample, while the bottom right shows examples of APs from different parts of the sample (shown by the multicolored rectangles in the map on the upper left).

To elucidate the reason for the heterogeneity of the AP, we simulated wave conduction over the same sample with different stimulus location (I_stim_). Figure 3A shows a comparison of stimulation along the upper edge (right) and the upper right corner of the sample: the S maps differ significantly for these two cases, i.e., the action potential of the same cell depends on the direction of the wave front passing through it (Figure 3B). Thus, changing the direction of the wave front changes the shape of the action potential both locally (Figure 3B) and globally (Figure 3C), changing the degree of AP heterogeneity within the sample: stimulation of the upper edge leads to a narrower distribution of the S value in the sample. The reason for the heterogeneity of the AP can be explained as follows, the AP of each cell depends on the local drain and source ratio: the number of gap contacts on different sides of the cell may not coincide, in which case the charge entering the cell and the charge needed to excite the next cells may differ, which determines the shape of the action potential. Wave propagation in the opposite direction reverses the drain-to-source ratio. Ephatic coupling between cells is possible across the membrane, not just at the cell ends, so the change in AP occurs with any shift in the stimulation angle. Our model showed that the heterogeneity of APs can be determined by tissue morphology and not only with different electrophysiology (expression of genes encoding ion channel proteins) [27].

**Figure 3.**
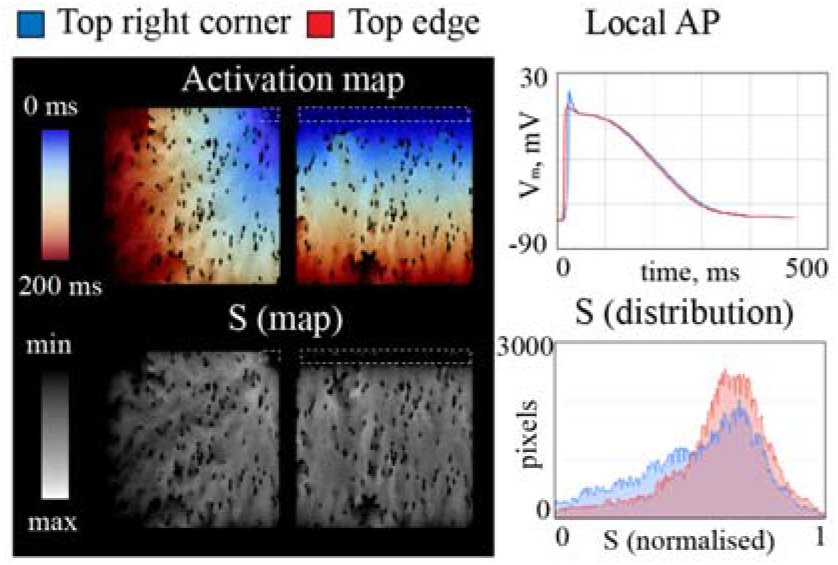
Activation maps (top) and AP morphology maps (bottom, black and white) for the same sample with different location of stimulation sites: upper right corner (left column) and upper edge of the sample (right column). The graph on the right shows the action potential in the same region (cell), with AP in red when the sample is stimulated at the upper edge, and in blue when the upper right corner is stimulated. Histograms of S-values with maps of AP morphology are presented on the lower right, the narrower distribution (red) corresponds to stimulation along the upper edge.

In addition, the distribution of the AP shape in space (Figure 3A, bottom) shows the presence of “conduction pathways” - dedicated directions within which the shape of the AP changes to a lesser extent. In Figure 3, the conductive pathways are directed from the stimulation point and bypass fibrous areas. Previously, the observation of conductive pathways was made in the case of high concentrations of FB [9]. The combination of conductive pathways and association of AP with the direction of the wave actually makes possible the occurrence of alternans depending on the tissue morphology, and not only on electrophysiology [28].

### 3.2. Simulation of spiral waves in heterogeneous tissue

One consequence of the heterogeneity of the AP is the possibility of forming a unidirectional block. Figure 4A shows a unidirectional conduction block based on the “diode” principle [29], when wave propagation is possible only in one direction because of the narrowing section of the conducting tissue (limited by fibroblasts). Figure 4B shows a spiral wave launched in a sample without fibroblasts: here the inhomogeneity of the AP was significant enough to create a unidirectional conduction block. In this case, the block leads to the stopping of the spiral rotation. Despite the absence of fibroblasts, the leading edge of the wave remains curved because of the inhomogeneity of the AP associated with the local drain and source ratios in the cells.

**Figure 4.**
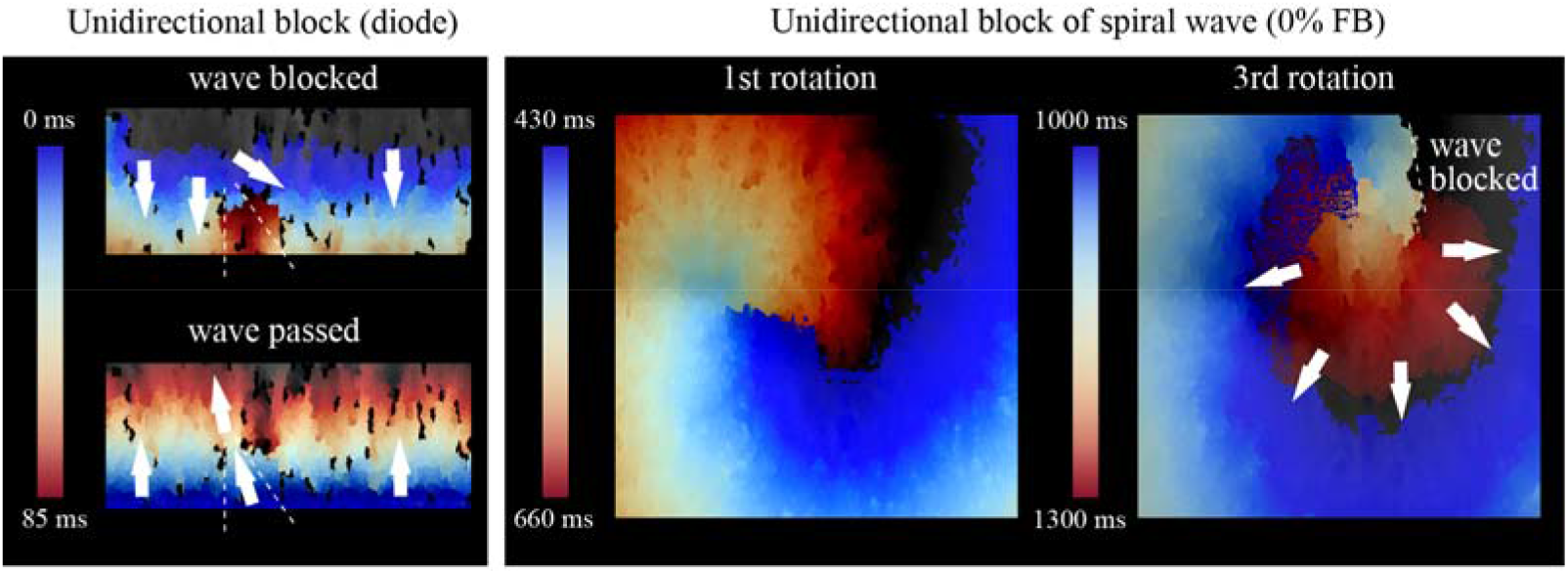
Left - unidirectional conduction block when the sample is stimulated from above. Stimulus can pass through the same region when the sample is stimulated from below. On the right - propagation of the spiral wave front in the sample without FB, occurrence of unidirectional block.

The addition of fibroblasts (10%) leads to a change in the trajectory of the spiral wave: the inhomogeneity of the AP here also causes the spiral to move along the sample, until the spiral core attaches to a zone with an increased concentration of FBs (yellow circle in Figure 5). In our model, both helix drift, which depends among other things on the different excitability [30] of the cells, and pinning to non-conductive areas are possible.

Thus, unidirectional block is possible in both cases (without FB and with FB), but fibroblasts are required for helix pinning. Having this distinction is an advantage of our discrete approach to modeling fibrosis.

**Figure 5.**
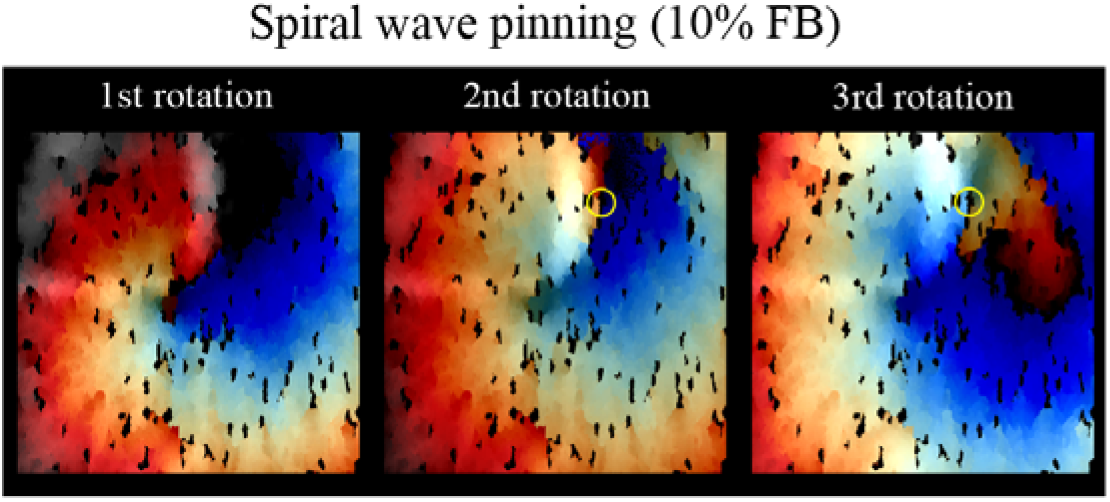
Spiral wave propagation trajectory in a sample with 10% FB, attachment of the spiral to a non-conductive region (yellow circle).

Figure 6 shows a comparison of spiral wave trajectories in a homogeneous sample (all diffusion coefficients D are equal), and in the layers generated by the Potts model (without FB and with 10%FB). The electrophysiology from the model [16] used to simulate fibrotic sites was used to generate a spiral in the homogeneous sample [16]. Comparing our proposed approach to modeling fibrosis with the already existing ones [16, 31], we can draw the following conclusions: unlike the approach [31] with randomly located non-conductive pixels, our model allows for AP inhomogeneity in the absence of fibroblasts and local sample anisotropy, which may not coincide with the global one set by the fiber or diffusion tensor direction. Our approach is also based on the random mixing of FBs and CMs in diffuse fibrosis, but the final shape of the cells is shaped by the Potts model and tends to form conductive pathways leading to a higher percolation threshold [9]. Modeling fibrosis as a homogeneous site [16] (no discrete FBs) leads to the impossibility of pinning the wave and curvature of the wavefront.

**Figure 6.**
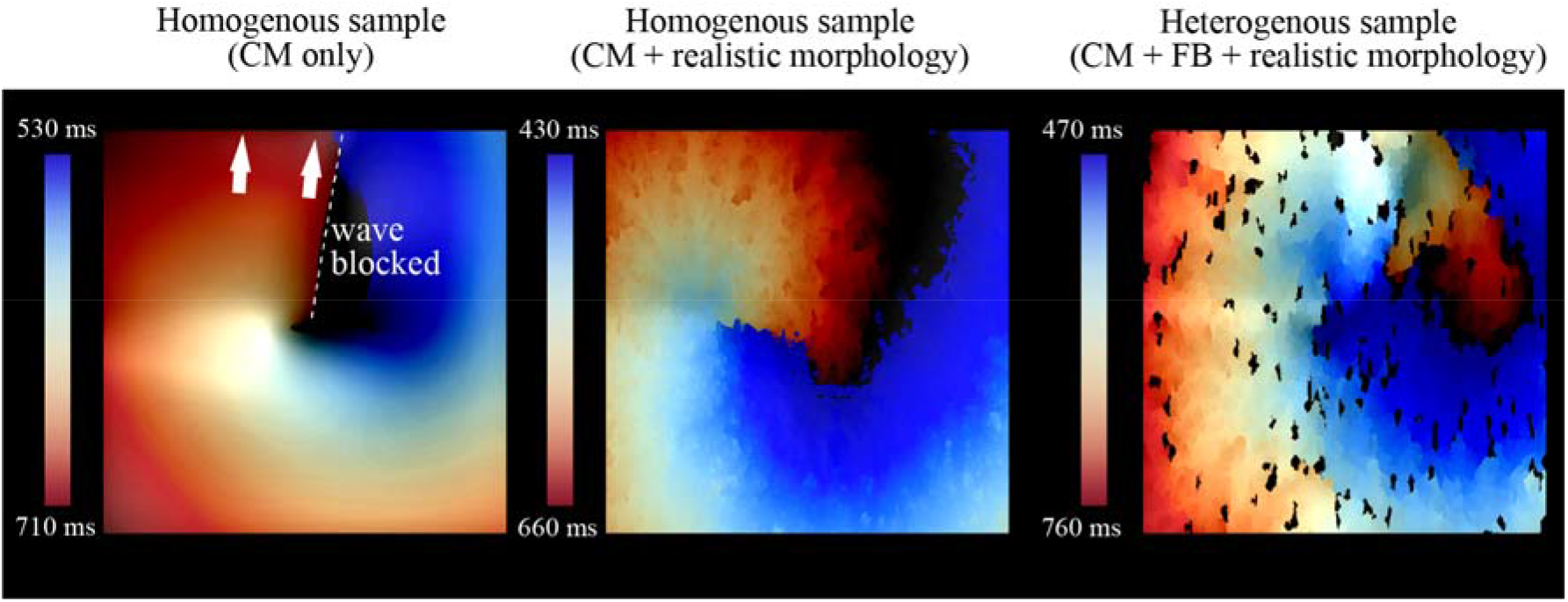
Comparison of spiral waves in a homogeneous sample (left), a sample without FB (middle), and a sample with 10% FB (right).

## 4. Conclusions

In this work, a computer model was developed and tested, which allows generating conductive tissue from cardiac cells with specified characteristics. The presented model explains the mechanism of connection of the AP shape with the tissue morphology with unchanged formulation of ionic currents. The relationship between the morphology of individual cells generated by the Potts model and the behavior of the excitation wave is shown, as well as the manifestation of nonlinear effects both at the level of individual cells (unidirectional block, heterogeneity of electrophysiology) and at the tissue level (meandering and pinning of the spiral wave). The key point is that the manifestation of these effects is not initially put in the model, but is a consequence of taking into account the morphology of cells and intercellular contacts, thus showing their role in ensuring a stable excitation conduction in the heart.

## Statements and Declarations

The authors declare no competing interests.

## Acknowledgments

We thank Dr. Kudryashova N.N. for her help with the Potts computer model. The work was supported by the strategic academic leadership program ‘Priority 2030’ (Agreement 075-02-2021-1316 30.09.2021) in Moscow Institute of Physics and Technology. We also thank «Tatneft» company for supporting the project.

